# Are these cardiomyocytes? Protocol development reveals impact of sample preparation on the accuracy of identifying cardiomyocytes by flow cytometry

**DOI:** 10.1101/388926

**Authors:** Matthew Waas, Ranjuna Weerasekera, Erin M. Kropp, Marisol Romero-Tejeda, Ellen Poon, Kenneth R. Boheler, Paul W. Burridge, Rebekah L. Gundry

## Abstract

Modern differentiation protocols enable efficient, yet imperfect, differentiation of human pluripotent stem cells into cardiomyocytes (hPSC-CM). As the number of laboratories and studies implementing this technology expands, the accurate assessment of cell identity in differentiation cultures is paramount to well-defined studies that can be replicated among laboratories. While flow cytometry is apt for routine assessment, a standardized protocol for assessing cardiomyocyte identity in hPSC-CM cultures has not yet been established. To address this gap, the current study leveraged targeted mass spectrometry to confirm the presence of troponin proteins in hPSC-CM and systematically evaluated multiple anti-troponin antibodies and sample preparation protocols for their suitability in assessing cardiomyocyte identity. Results demonstrate challenges of interpreting data generated by published methods and informed the development of a robust protocol for routine assessment of hPSC-CM. Overall, the new data, workflow for evaluating fit-for-purpose use of antibodies, and standardized protocol described here should benefit investigators new to this field as well as those with expertise in hPSC-CM differentiation.

## Introduction

Directed differentiation of human pluripotent stem cells (hPSC) into cardiomyocytes (hPSC-CM) offers an inexhaustible supply of cells for basic science research and translational applications, including drug testing, disease modeling, and regenerative medicine. Using modern differentiation protocols, hPSC-CM can be efficiently generated from human embryonic (hESC) and induced pluripotent stem (hiPSC) cells, which has led to an increasing number of laboratories and studies implementing this technology (reviewed in (Batalov and Feinberg, 2015; Mummery et al., 2012). However, despite significant advancements in defining the factors most critical for cardiomyogenic differentiation (Burridge et al., 2014; Lian et al., 2015), the resulting cultures remain a heterogeneous mixture with regards to cell type, maturation stage, and subtype, and this heterogeneity can be exacerbated by variations among cell lines, protocols, and personnel (Ohno et al., 2013). Ultimately, as heterogeneity can pose challenges to interpreting functional data, the ability to accurately and precisely assess cell identity in differentiation cultures is paramount to well-defined and reproducible studies.

Flow cytometry is a quantitative, cell population-based single-cell approach to assess individual cell phenotypes, rendering it an ideal strategy for assessment of hPSC-CM heterogeneity. In this approach, population heterogeneity is typically assessed based on detection of endogenous proteins by specific monoclonal antibodies or expression of exogenous marker proteins driven by a cell- or tissue-restricted promoters. Considering the availability of benchtop cytometers and prevalence of flow cytometry core facilities at most research organizations, this approach is affordable and accessible to most laboratories. Altogether, flow cytometry is well-suited for use in routine quality control assessments of hPSC-CM cultures. The proper implementation of flow cytometry requires optimization of many procedural parameters within sample preparation, data acquisition, and data analysis. Examples include optimizing the cell collection method to produce single cell suspensions, validating monoclonal antibody specificity, titrating antibody concentrations, selecting appropriate negative and positive controls, adjusting cytometer laser settings, and developing acceptable gating strategies. Considering the numerous procedural variables, this optimization process can be daunting. Unfortunately, a standardized and validated protocol that is broadly applicable among laboratories has not been established for assessing cardiomyocyte identity within hPSC-CM cultures. Consequently, accurate comparisons of outcomes generated by various differentiation protocols or cell lines among laboratories and studies, including assessments of purity, reproducibility, and functional data, remain challenging.

A survey of studies published over the past seven years (1/2010 - 10/2017) reveals a wide range of antibodies and experimental conditions reported for flow cytometry-based assessment of hPSC-CM. Of the 84 studies that use flow cytometry, the majority (n = 68) used cardiac troponin T (TNNT2) as the primary marker to assess hPSC-CM cultures (Figure 1A). Of these studies, nearly 72% used one of two monoclonal antibodies (clones 13-11 or 1C11), and 28% used a variety of other antibodies, including monoclonal and polyclonal, to detect TNNT2. Of concern, 18% of TNNT2 studies failed to report either the antibody clone, the vendor, or both. The sample preparation conditions among studies were more disparate (Figure 1B), with nine fixation and fifteen permeabilization conditions reported. Moreover, many studies and failed to report the relevant details for fixation (>15%) and permeabilization (>26%). Altogether, there is currently no consensus regarding which marker, antibody, or protocol is best suited to enable comparisons of hPSC-CM culture heterogeneity among experiments or laboratories.

**Figure 1.**
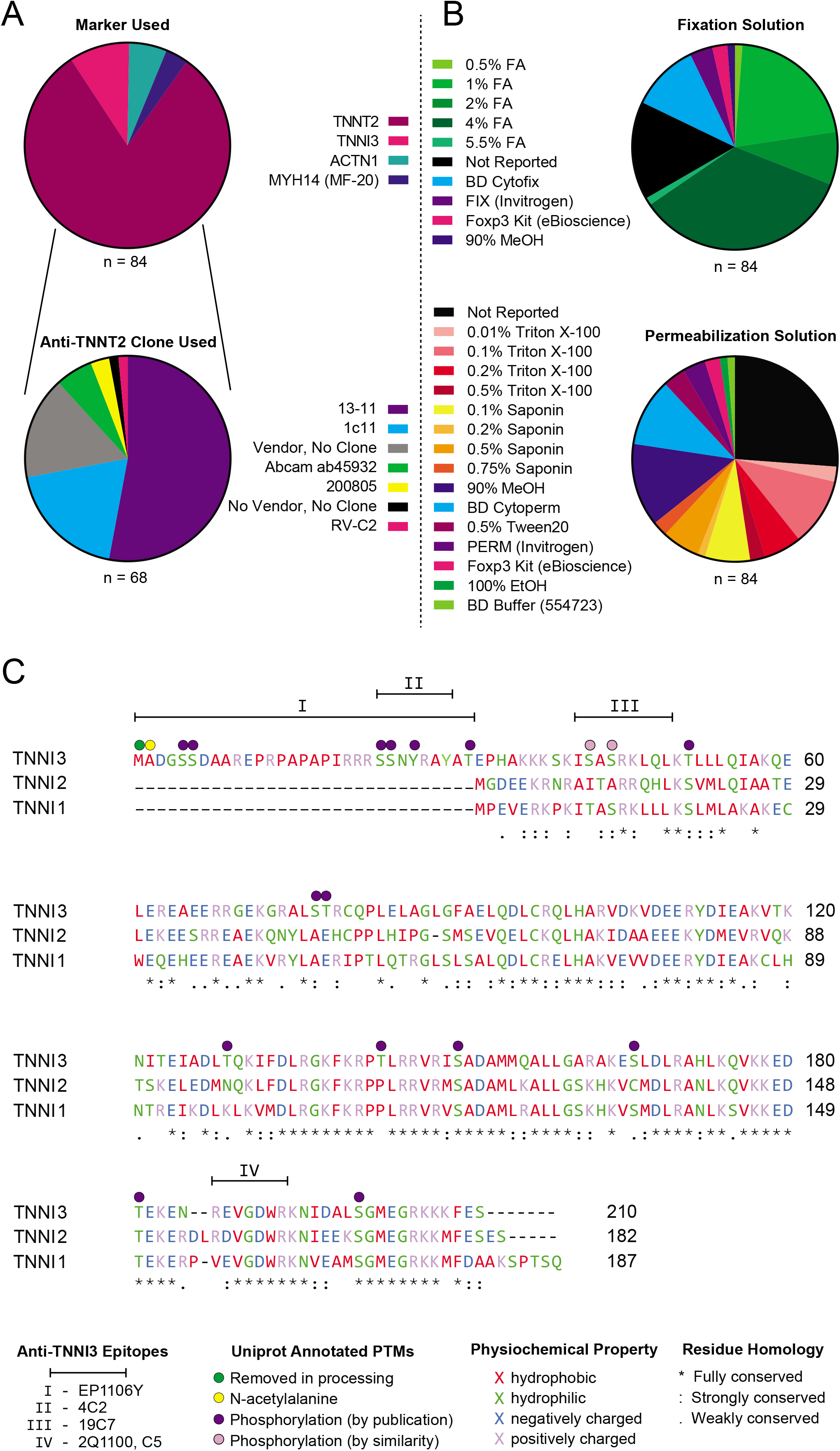
Results from the literature survey of antibodies and sample preparation techniques used for flow cytometry assessment of hPSC-CM cultures. (A) Chart summarizing the protein markers used, and expanded chart showing the variety of antibodies used to detect TNNT2. (B) Chart summarizing sample fixation and permeabilization methods used for any of the antibodies in literature summarized in panel A. (C) Alignment of troponin I isoforms showing alignment information, post-translational modifications, and the reported epitopes for anti-TNNI3 monoclonal antibodies evaluated here.

The troponin family is the protein class most commonly used for determining cardiomyocyte identity in hPSC-CM cultures (Figure 1A). The troponin complex is located on the thin filaments of striated muscle cells and is responsible for regulating contraction (Ebashi, 1983). This complex is composed of three protein subunits - troponin T, troponin C, and troponin I. Each troponin gene gives rise to multiple protein isoforms (TNNT1-3; TNNC1-3; TNNI1-3) whose expression are relatively restricted among muscle types (*i.e*. cardiac, smooth, slow skeletal, and fast skeletal) (Sheng and Jin, 2016). While TNNT2 has been the most commonly used marker for hPSC-CM assessment by flow cytometry in the studies surveyed here, TNNT2 has been detected in skeletal muscle (Anderson et al., 1991; Bodor et al., 1997) and multiple types of smooth muscle (Bicer and Reiser, 2013; Kajioka et al., 2012). In contrast, TNNI3 expression is restricted to the cardiomyocyte throughout human development (Bodor et al., 1995; Rittoo et al., 2014).

Considering the lack of consensus regarding marker, antibody, and protocol, the broad goals of this study were to evaluate antibody specificity and sample preparation conditions for the assessment of cardiomyocyte identity within hPSC-CM cultures by flow cytometry. Three sample preparation methods in conjunction with five commercially available anti-TNNI3 and two anti-TNNT2 antibodies were applied to hPSC-CM and two negative control cell types, undifferentiated hPSC and cardiac fibroblasts. In performing these analyses, we found that the choice of fixation protocol and antibody had significant and variable effects on the accuracy of cardiomyocyte identity assessment. It is expected that by providing details regarding validation of antibody specificity within this context and revealing pitfalls with commonly used antibodies and preparation conditions, these results will benefit laboratories with established expertise in hPSC-CM differentiation as well as those new to this field. By establishing rigorous standards for quality control evaluation of hPSC-CM, we believe that the approaches described here will facilitate the use of hPSC-CM in a broad range of research and clinical applications, especially by enabling more accurate comparisons of results among studies. To facilitate data sharing among laboratories, the current study aims to set a standard regarding the experimental details that should be included when publishing flow cytometry-based assessments of hPSC-CM, consistent with similar calls for publication guidelines (Lee et al., 2008). Finally, based on results of the current study, a comprehensive protocol for assessment of cardiomyocyte identity in hPSC-CM cultures by flow cytometry is provided. The protocol provides stepwise instructions and describes key points to consider for sample preparation and antibody validation, with the expectation that providing these details will facilitate its use among laboratories. As this protocol has been successfully replicated in three different laboratories and can be completed, from adherent-cell collection to data analysis, in less than three hours, it is suitable for routine assessment of hPSC-CM cultures.

## Results

### Targeted Mass Spectrometry for Detecting TNNI3 and TNNT2 in hPSC-CM

The expression of troponin complex components is temporally regulated during normal human development in a tissue-specific manner (Bhavsar et al., 1991; Hunkeler et al., 1991; Sasse et al., 1993). While similar trends in temporal regulation have been reported for *in vitro* differentiation of hPSC-CM, discrepancy with regards to the timing of the emergence of TNNI3 has been reported, with one report suggesting it emerges after months in culture (Bedada et al., 2014) and others showing it is expressed as early as day 8 (Puppala et al., 2013; Tompkins et al., 2016). For this reason, we used a targeted mass spectrometry approach to confirm the presence of TNNI3 and TNNT2 protein in day 25 hPSC-CM as a first step in the selection of reliable markers of cardiomyocyte identity. The approach, parallel reaction monitoring (PRM), uses high resolution/accurate mass instrumentation to specifically detect pre-selected peptides within a mixture (Peterson et al., 2012). Here, PRM assays were developed to specifically detect three unique peptides from TNNI3 and four from TNNT2. Stable isotopically labeled peptides for TNNI3 were included as internal controls to provide added rigor for this protein because of reported discrepancies regarding timing of its expression. Application of this PRM assay reliably detected peptides from both TNNI3 and TNNT2 in day 25 hPSC-CM and in human cardiac tissue, but not in undifferentiated hPSC (Figure 2, Figure S1). Importantly, for TNNI3 peptides, the endogenous and isotopically labeled peptides co-eluted and had identical fragmentation patterns across hPSC-CM and cardiac tissue, providing unequivocal evidence that this protein is present in these samples (Figure 2). Altogether, this highly sensitive, antibody-independent mass spectrometry strategy confirms the presence of TNNI3 and TNNT2 in day 25 hPSC-CM produced by the differentiation protocol used here.

**Figure 2.**
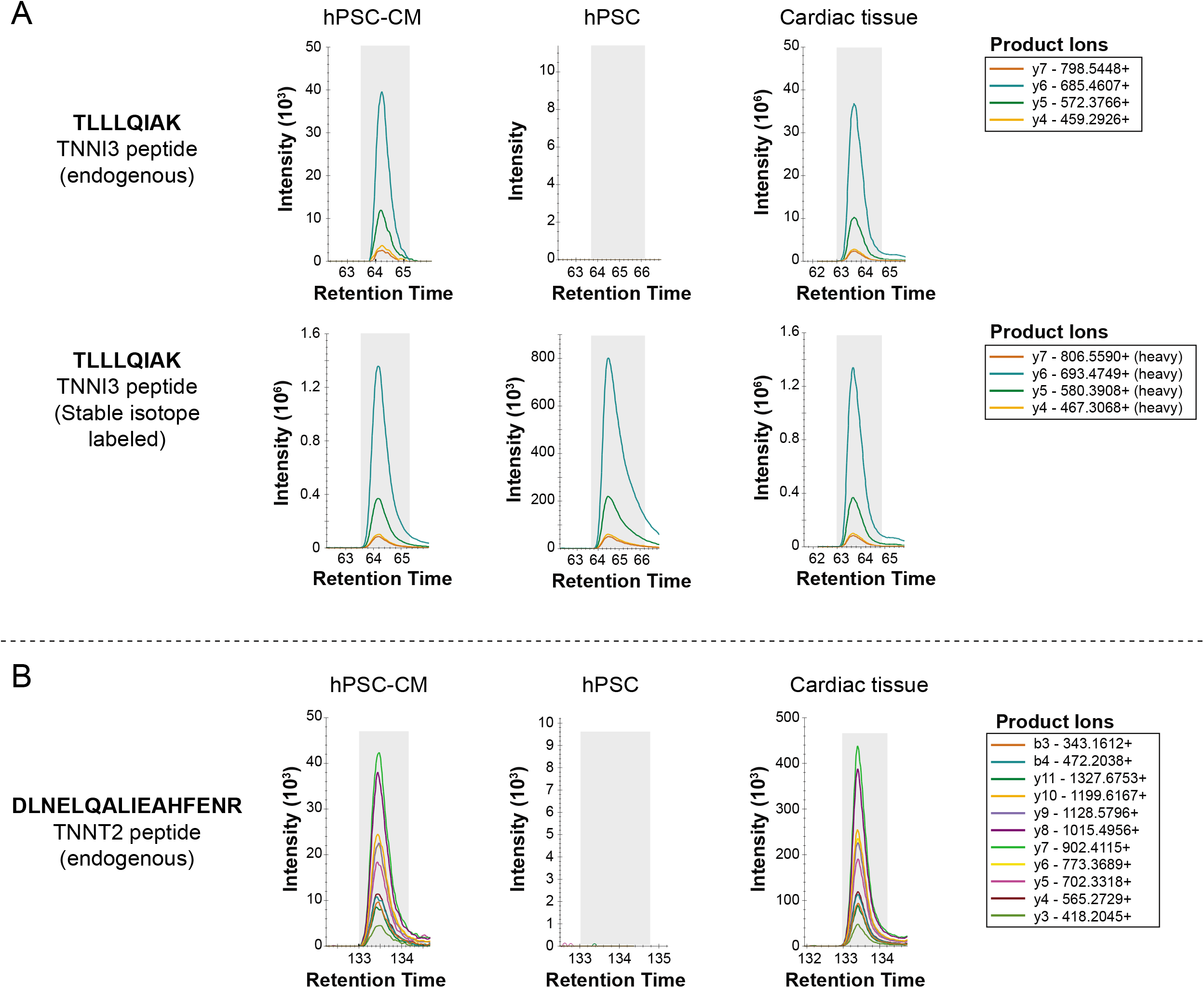
Targeted mass spectrometry results for TNNI3 and TNNT2 in hPSC-CM, hPSC, and cardiac tissue. (A) Extracted ion chromatograms for all monitored product ions belonging to an endogenous TNNI3 peptide and its stable isotope labeled (SIL) control peptide in hPSC-CM, hPSC, and cardiac tissue. Product ions for endogenous and SIL peptides co-elute and the peak areas for corresponding product ions are proportional to each other. (B) Extracted ion chromatograms for all monitored product ions belonging to an endogenous TNNT2 peptide in hPSC-CM, hPSC, and cardiac tissue. Peak areas for corresponding product ions between hPSC-CM and cardiac tissue are proportional to each other. The assigned peak boundaries for all peptides are designated by the gray shaded area. Data for additional peptides belonging to TNNI3 and TNNT2 are shown in Figure S1.

### Antibody Clone and Sample Preparation Screen

Although our literature survey revealed that a preponderance of studies relied on TNNT2 as a marker of cardiomyocyte identity, TNNI3 is more specific to cardiomyocytes than TNNT2 throughout human development. As the mass spectrometry analysis confirmed the presence of both proteins in day 25 hPSC-CM, antibodies to both TNNI3 and TNNT2 were investigated for their ability to serve as markers of cardiomyocyte identity within hPSC-CM cultures. The two most common troponin T antibodies from previous studies (Figure 1A) and five commercially available anti-TNNI3 antibodies whose epitopes span the range of the amino acid sequence for TNNI3 (Figure 1C) were assessed for their ability to specifically detect hPSC-CM using three different sample preparation conditions (Table 1). In the initial screen, all seven antibodies were assessed for their ability to produce signal stronger than that of an equivalent amount of isotype control and to distinguish hPSC-CM from undifferentiated hPSC, a relevant negative cell-type control. All data for two biological replicate analyses of each clone and sample preparation protocol are presented in Figure S2A-G. Overall, flow cytometry results were highly dependent on sample preparation conditions for some antibodies, and less-so for other clones (summarized in Figure 3A, supporting data in Figure S2). For example, all three sample preparation protocols yielded satisfactory results for clone 1C11, but the ability to distinguish between positive and negative cell types was protocol-dependent for clones 13-11 and 2Q1100 (Figure 3B). Clones 19C7 failed to produce desirable results independent of protocol as it produced a stronger signal in the negative cell type control than in hPSC-CM (Figure 3B). Each sample preparation strategy can produce samples suitable for flow cytometry demonstrated by single-cell suspensions that were separable from debris and dead cells by gating on forward and side scatter (Figure S2) and scatterplots for all subsequent experiments were comparable with those shown in Figure S2. However, samples prepared using protocol 2 exhibited more favorable handling characteristics (*i.e*. a tight, visible cell pellet) and, in general, better resolution compared to protocols 1 and 3. Consequently, protocol 2 and the four antibodies (1C11, 13-11, C5, 2Q1100) which provided the most satisfactory results during the initial screen were assessed further in subsequent experiments.

**Figure 3.**
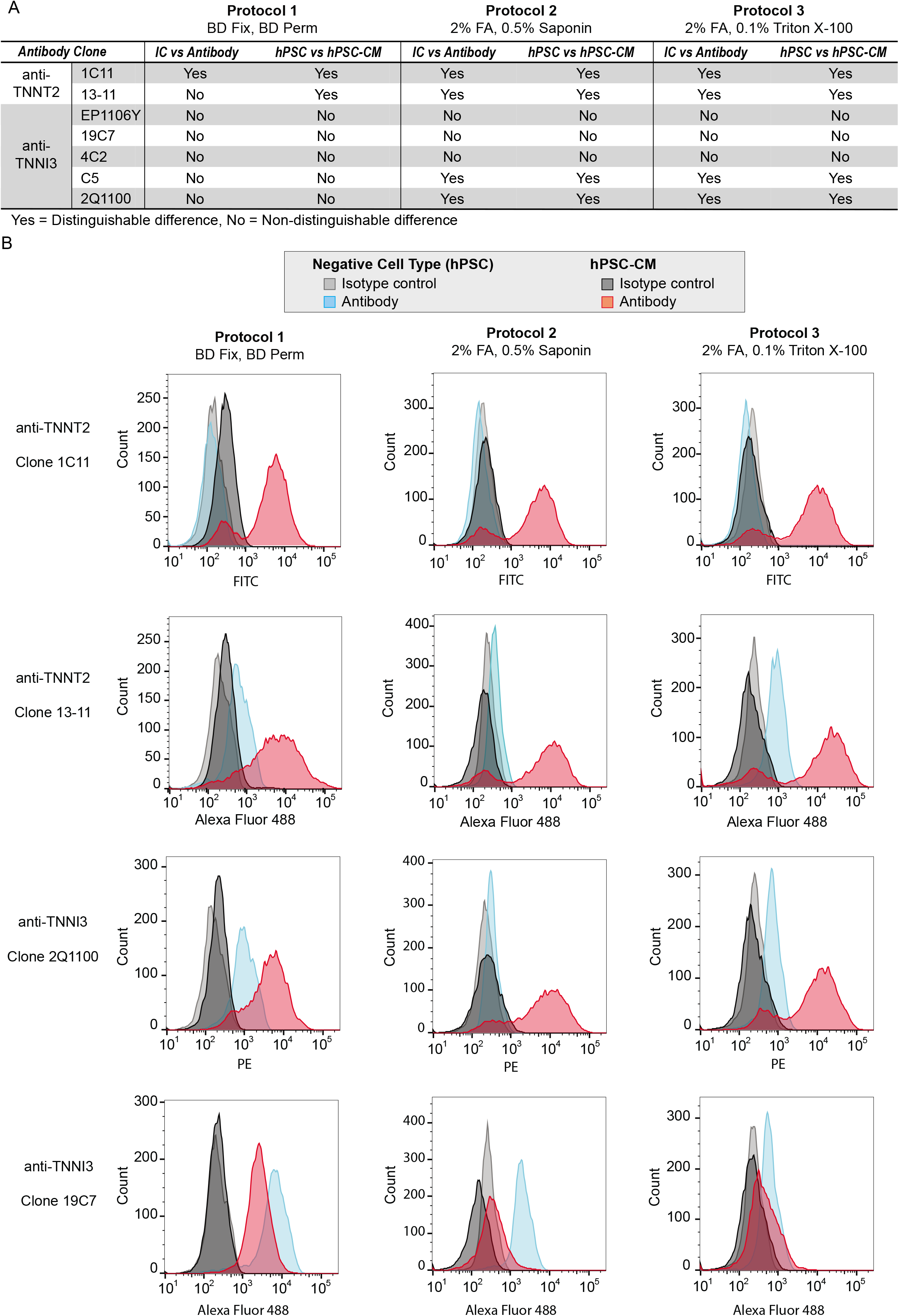
Results of the initial screen of seven antibodies using the three sample preparation protocols applied to hPSC-CM and hPSC. (A) Summary of whether antibodies provided acceptable results (*i.e*. positive signal for hPSC-CM and negligible signal for hPSC; positive signal for antibody with negligible signal for isotype control (IC)) or unacceptable results (*i.e*. non-specific binding to hPSC, insufficient signal for hPSC-CM). (B) Histograms for selected antibodies demonstrating the range of effects the sample preparation had among antibodies. Histograms for all data are provided in Figure S2A-F.

**Table 1.**
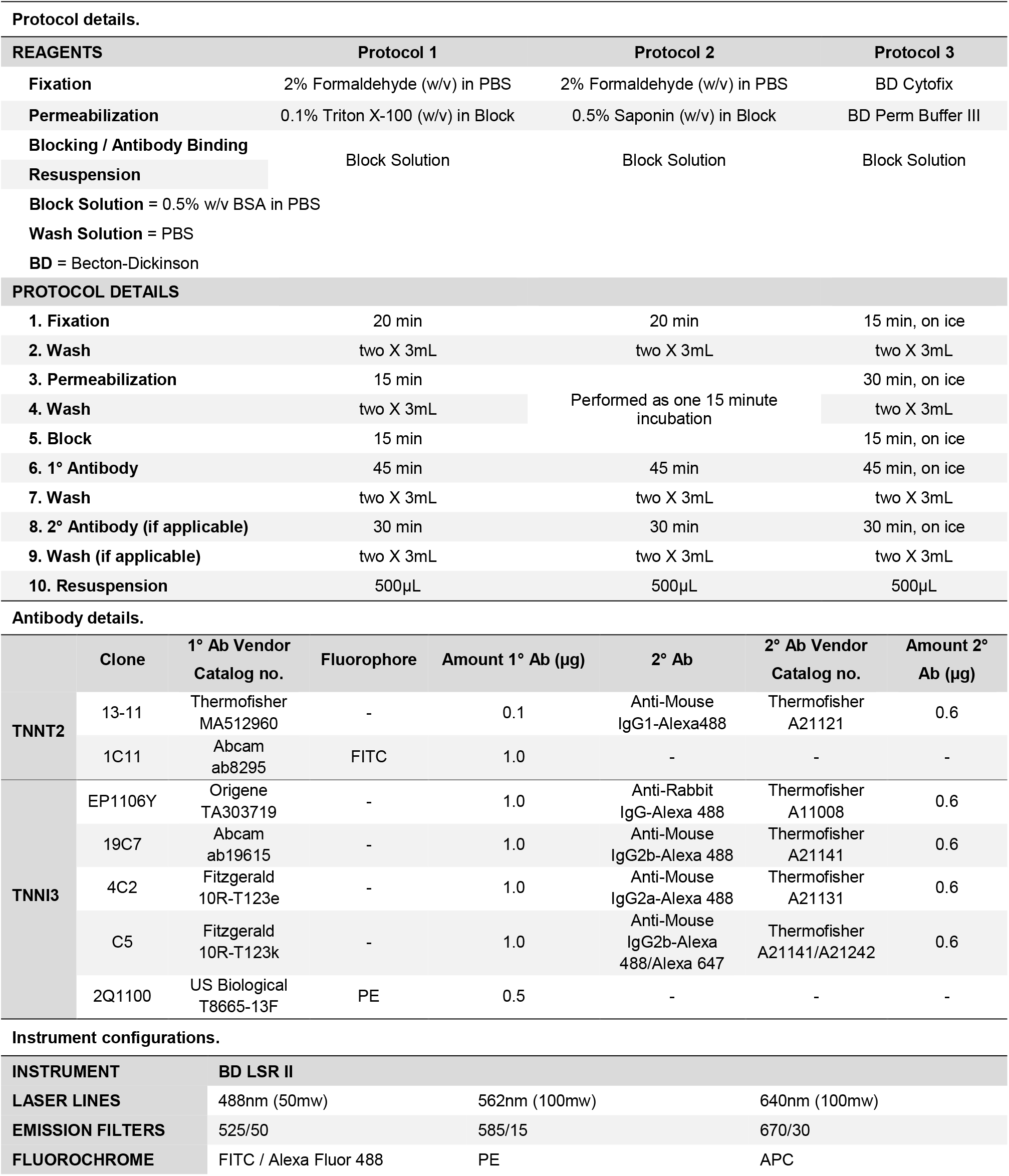
Summary of the experimental conditions examined for their suitability for assessing hPSC-CM cultures by flow cytometry. Details are provided for the three sample preparation protocols (A), the seven antibodies evaluated (B), and the flow cytometer instrument configurations (C).

### Antibody Titration

Four concentrations were tested for each antibody, based either on vendor recommendations or the results of the screen, to determine the optimal concentration for providing a maximal separation in signal between positive and negative cell types (Figure 4A, Figure S3). Performance of all four antibodies was consistent with results expected for a successful titration study (*i.e*. signal dependent on antibody concentration that eventually becomes saturated in a positive population) (Figure 4A, S3). Three clones (1C11, 2Q1100 and C5) that were best able to distinguish between negative and positive populations were selected for further validation using the optimal antibody amount determined by titration – 0.5 μg for 1C11 and 2Q1100, 0.1 μg for C5.

**Figure 4.**
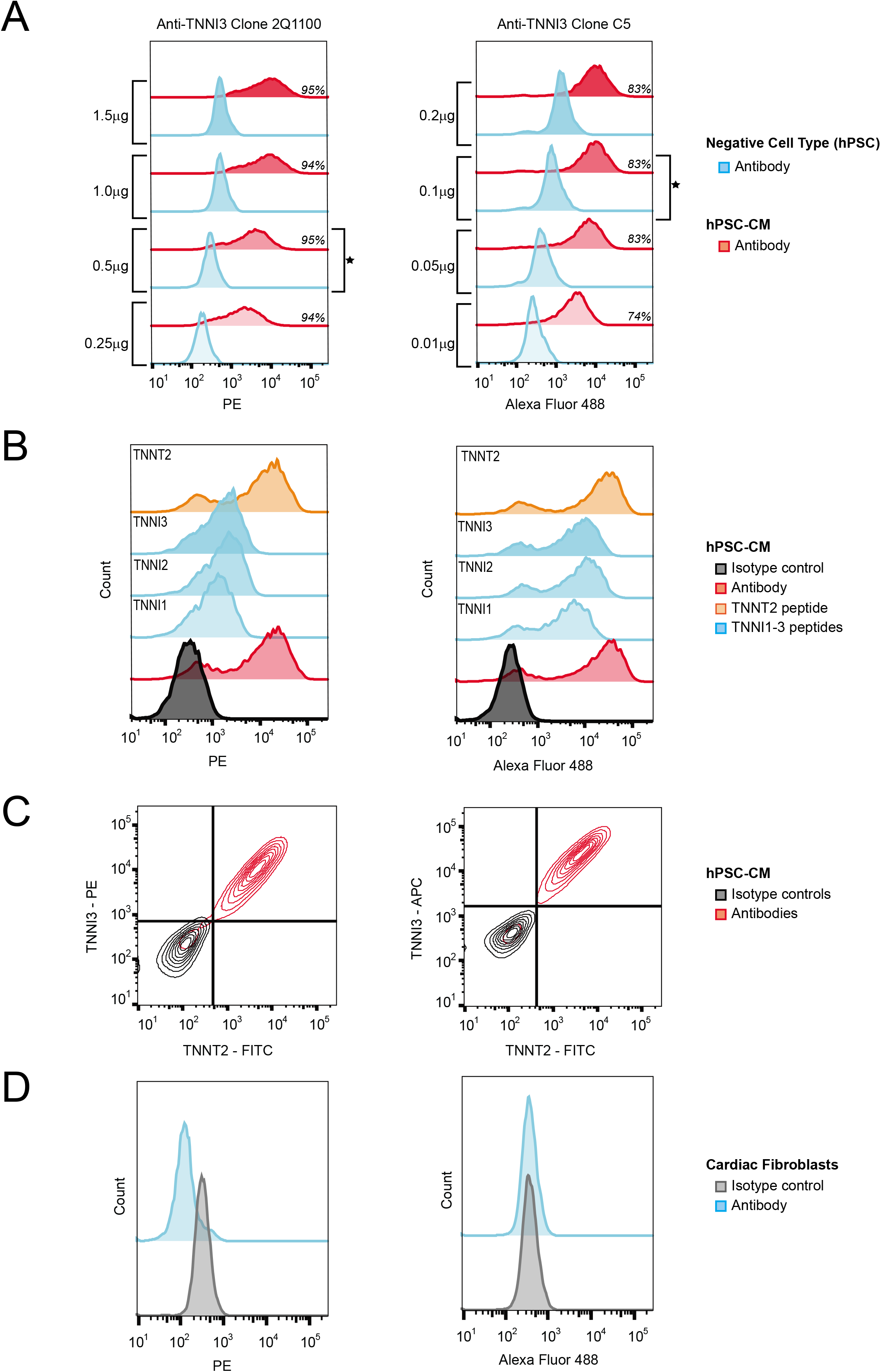
Results from antibody characterization experiments for anti-TNNI3 clone 2Q1100 and anti-TNNI3 clone C5. (A) Antibody titration histograms demonstrating saturable signal with the percentage of positive cells for each amount of antibody in italics and the selected concentration marked by a star. (B) Epitope competition assay histograms labeled with the peptide epitopes pre-incubated with the antibodies depicting the partial blocking by TNNI peptides of anti-TNNI3 clone 2Q1100 but not anti-TNNI3 clone C5. (C) Co-immunodetection contour plots showing that anti-TNNT2 clone 1C11 and anti-TNNI3 clones are marking the same population of cells with the hPSC-CM culture. (D) Histograms from cardiac fibroblast control experiments signifying no binding of anti-TNNI3 clones. All experiments used sample protocol 2. Data from all replicates (*n* = 3 for each) are shown in Figure S3-S4.

### Epitope Competition Assay, Co-Immunodetection, and an Additional Negative Cell Type Control

The specificity of clones 1C11, 2Q1100 and C5 for their reported epitopes was assessed using a competition assay in which signal from each naïve antibody was compared to antibody preincubated with peptide antigen. In this manner, a diminution or ablation of signal caused by incubation with peptide antigen can be indicative of specificity for the reported epitope. Due to the high sequence identity between the isoforms of TNNI1, TNNI2, and TNNI3 at the reported epitope for both clones C5 and 2Q1100, the homologous peptide regions to the TNNI3 epitope were also investigated. Antibody incubated with TNNT2 and TNNI3 epitopes were included as negative controls for anti-TNNI3 and anti-TNNT2, respectively, for these experiments. Amino acid sequences for purified peptides antigens are shown in Figure 1C. TNNI1, TNNI2 and TNNI3 peptides were able to partially block anti-TNNI3 clone 2Q1100 binding to hPSC-CM shown by the overall decrease in fluorescence intensity and collapse of the histogram into a unimodal distribution. In contrast, these peptides only had a minor effect on binding of anti-TNNI3 clone C5 to hPSC-CM (Figure 4B, Figure S4A). Compared to naïve antibody, anti-TNNT2 clone 1C11 antigen peptide had no effect on the clone 1C11 histogram (Figure S4A). Although the epitope competition assay was unable to unequivocally verify specificity of the antibodies for their reported peptide epitopes, this may be simply due to a linear peptide lacking the necessary secondary or tertiary structure of the native epitope. Consequently, a co-immunodetection strategy was used to determine if antibodies to TNNI3 and TNNT2 were specific to the same cell population as an alternative assessment of specificity. If they do not overlap, this would suggest that one or both is binding to non-cardiomyocytes. When hPSC-CM were evaluated with anti-TNNI3 and anti-TNNT2 antibodies using the following clone pairs: 2Q1100/1C11 and C5/1C11, less than 3% of the population, on average, was positive only for a single antibody *(i.e*. 97% of cells were positive for both antibodies or for neither). These results demonstrate that, under these preparation conditions, the TNNI3 and TNNT2 antibodies used here mark the same cell population (Figure 4C, Figure S4B). These results, together with the observation that immunofluorescent imaging experiments using anti-TNNT2 clone 1C11 yield a striated localization pattern expected for a sarcomere protein (Figure S4D), support that these antibodies are specifically detecting their respective protein targets when used with this sample preparation protocol. Finally, while the application of this protocol within the context of hPSC differentiation was the focus of this study, the protocol was also applied to cardiac fibroblasts, a biologically relevant negative cell type in co-culture (Thavandiran et al., 2013) and trans-differentiation experiments (Addis et al., 2013; Fu et al., 2013). Overall, using this protocol, all three antibody clones generated histograms from cardiac fibroblasts that were indistinguishable from isotype control (Figure 4D, S4C).

### Evaluating Protocol Performance in Mixed Populations and Among Laboratories

To accurately determine the percentage of cardiomyocytes within a heterogeneous hPSC-CM culture, a protocol, including antibody and all experimental conditions, must be able to distinguish cardiomyocytes from non-cardiomyocytes within a single tube. To evaluate the best performing protocol for this capacity, three antibody clones (1C11, 2Q1100 and C5) were used in conjunction with protocol 2 to assess population heterogeneity within samples where hPSC-CM and hPSC were mixed at various ratios (100:0, 75:25, 50:50, 25:75, and 0:100). Overall, each antibody clone in conjunction with sample preparation protocol 2 can distinguish between positive and negative cell types (Figure 5, Figure S5). At each ratio of hPSC-CM to hPSC, a bimodal population was observed where the percent positive cells decreased in proportion to the number of hPSC added to the sample (Figure 5, Figure S5). The percent positivity observed for samples that were a mix of hPSC-CM and hPSC correlated well with the expected percentages calculated based on the unmixed sample, averaging less than an 8% difference. Deviations from expected percentages are likely due to variations in cell counting as evidenced by dissimilarities in the event rates observed on the flow cytometer (data not shown) and by the increase in the percent errors that correlated with amount of hPSC added (Figure 5, S5). Considering the strong performance of these antibodies and protocol in cell mixing experiments, a detailed standard operating procedure was established and shared with two laboratories located in different institutions to further test rigor and reproducibility. Results from these two laboratories were comparable with our own data, despite using different cell lines and differentiation protocols, and similar trends were observed for correlations between expected and measured percent positivity and the maintenance of a bimodal population across samples (Figure 5).

**Figure 5.**
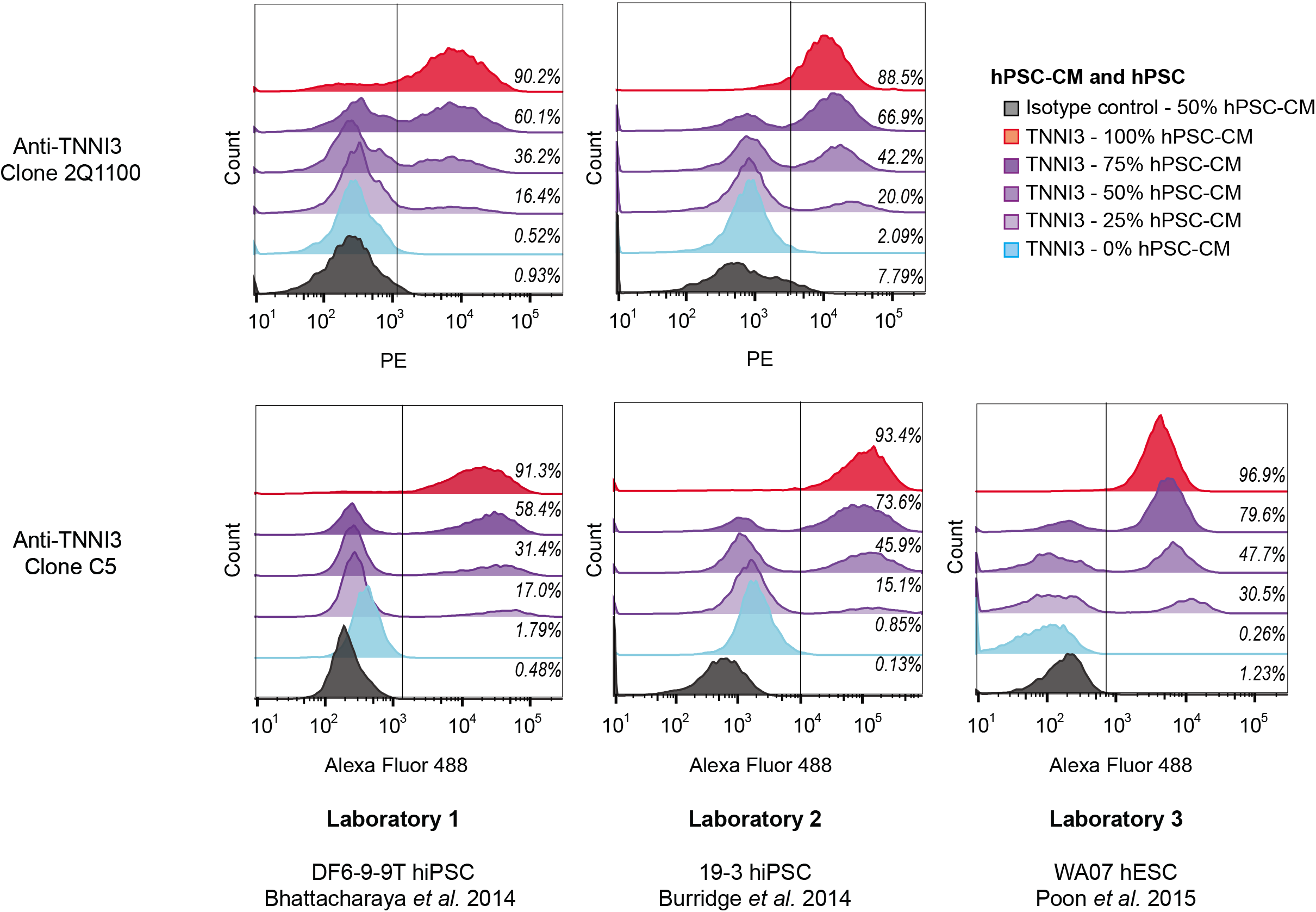
Results from mixed population experiments for anti-TNNI3 clone 2Q1100 and anti-TNNI3 clone C5. Histograms for the various populations are shown with the gates drawn and the percentage of positive cells for each condition listed in italics. For both anti-TNNI3 clones, the percent positivity decreased relative to the proportion of hPSC added to the sample. Data are consistent among laboratories, independent of the cell line and differentiation protocol (labeled under the histograms). All experiments used sample protocol 2. Data from all replicates (n = 3 for each) are shown in Figure S5.

### A Generalizable Workflow for Establishing Fit-for-Purpose Use of Antibodies

The spirit of this study is responsive to recent calls for improving scientific rigor and reproducibility discussed in several recent publications (Bordeaux et al., 2010; Bradbury and Pluckthun, 2015; Brooks and Lindsey, 2018) and reflected in policies for reagent validation that are now required by granting agencies (*e.g*. NIH). Our success in developing a replicable protocol supported the development of a standardized workflow for rigorous selection and evaluation of antibodies and sample preparation conditions for flow cytometry experiments (Figure 6). This workflow outlines major steps required to establish the fit-for-purpose of an antibody and protocol for assessing cell population identity within a heterogeneous mixture. To be clear, although data from vendors or previous publications can serve as starting points, antibody validation is ultimately the responsibility of the user and should be performed for each antibody clone, cell type, and protocol. To begin, suitable markers can be selected from literature or experimentally determined by using mass spectrometry. The superior selectivity, sensitivity, and specificity of targeted mass spectrometry make it an ideal technique for verifying the presence of candidate markers in cell types of interest compared to antibody-based techniques such as immunoblotting (Aebersold et al., 2013). The selection of antibody clones should consider published literature and vendor data as well as specific information about the epitope including uniqueness of the sequence and possible variants or post-translational modifications. As exemplified in this study, it is advisable to test more than one antibody clone and more than one protocol. Following antibody selection, screening, titration, specificity testing, and range-of-use demonstration should rigorously evaluate the utility of the antibody and protocol for its ability to specifically identify the cell type of interest within a heterogeneous sample. The overall approach and desired results for verification experiments are outlined in Figure 6. Based on this workflow and results from the current study, a comprehensive standard operating protocol (SOP) is provided in the supplemental data. The SOP describes these experiments in more detail and contains suggestions and considerations for the design of these experiments, and others, as well as their limitations.

**Figure 6.**
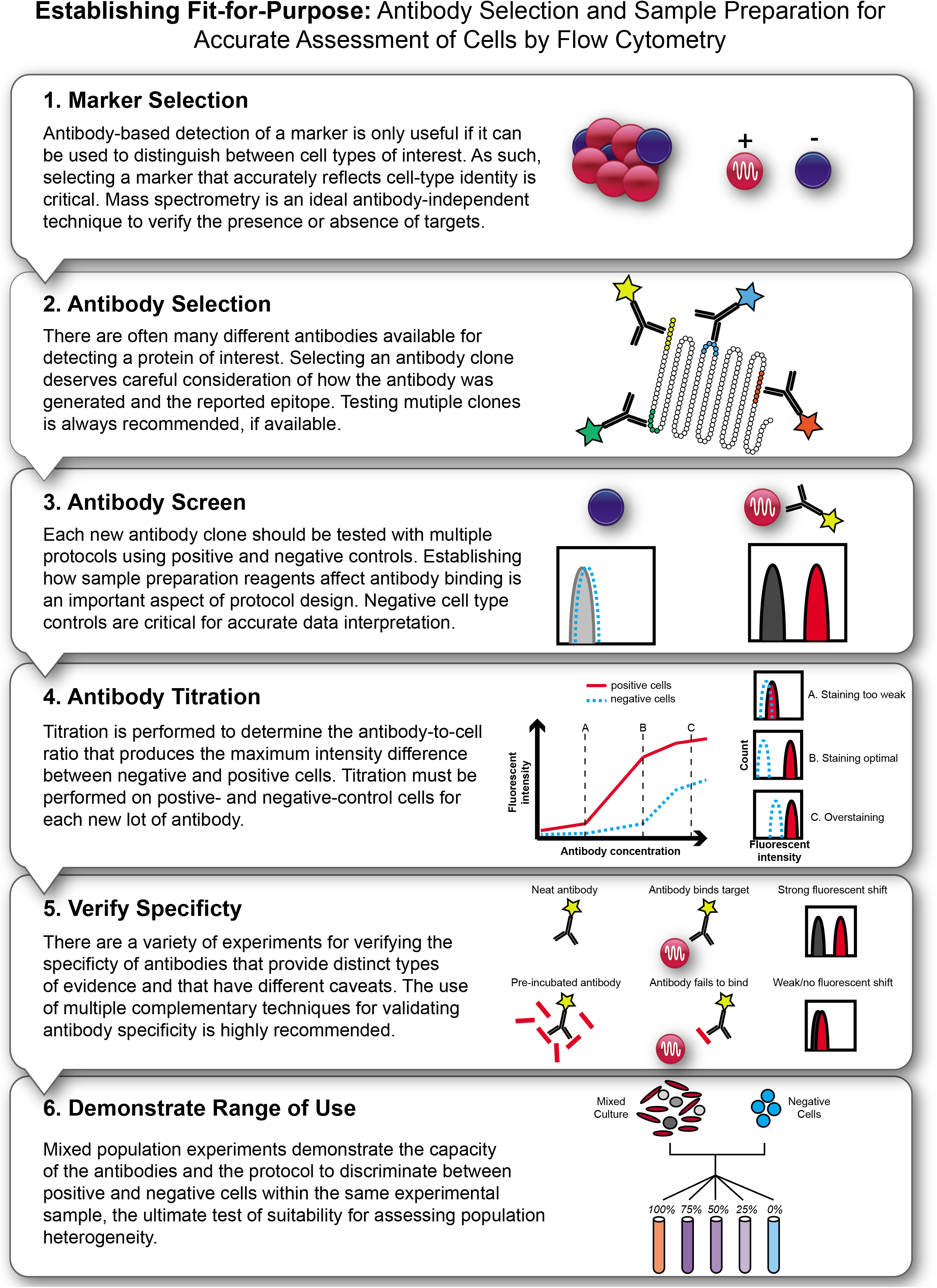
Workflow for establishing the fit-for-purpose of a flow cytometry protocol. The workflow outlines a stepwise progression through the steps necessary to establish the fit-for-purpose of antibodies and sample preparation protocols for flow cytometry experiments. Details and suggestions for experimental design are provided in the Supplemental Information as part of the standard operating procedure (SOP).

## Discussion

To date, a wide array of antibodies and sample preparation methods to assess hPSC-CM cultures have been described in published literature. Consequently, protocols for assessing the percentage of troponin-positive cells have not yet become standardized, posing challenges to comparing results among laboratories, differentiation protocols, cell lines and personnel. The aim of this study was to establish the fit-for-purpose of a flow cytometry protocol for assessing the percentage of cardiomyocytes within an hPSC-CM differentiation culture. Specifically, the study was designed to establish and subsequently validate that a protocol, (*i.e*. defined sample preparation, antibody clones, antibody concentrations), can perform its specified purpose (*i.e*. identify cardiomyocytes within hPSC-CM culture) to a specified level of quality (*i.e*. reliable and replicable). The results from these studies informed the generation of a SOP which includes relevant suggestions and observations to assist implementation of best practices in flow cytometry consistent with other recent calls for increased rigor with respect to antibody use (Bordeaux et al., 2010; Bradbury and Pluckthun, 2015; Brooks and Lindsey, 2018).

Two members of the intracellular troponin complex, TNNT2 and TNNI3, have been the most popular protein markers used in flow cytometry-based assessments of hPSC-CM. However, the utility of TNNT2 as a specific cardiomyocyte marker may be complicated due to its presence in various types of smooth muscle and in skeletal muscle during early development. While this concern might be insubstantial in the context of hPSC-CM differentiation where there is little evidence to indicate modern differentiation protocols routinely generate skeletal or smooth muscle, it remains a consideration. In contrast, TNNI3 is more specific to cardiac myocytes throughout development, yet this has been a less popular marker among the studies surveyed here. This may be due to convention, or, as demonstrated in this study, due in part to a lack of reliable antibodies. Another reason for avoiding TNNI3 as a marker of cardiomyocyte identity during hPSC-CM differentiation may be the uncertainty regarding the timing of its expression. To address this uncertainty, we used a targeted mass spectrometry approach to confirm that TNNI3 is present in day 25 hPSC-CM.

Proceeding with the knowledge that both TNNT2 and TNNI3 protein are present in day 25 hPSC-CM, we designed an antibody screen to test the suitability of three sample preparation protocols on positive and negative cell types for seven commercially available monoclonal antibodies reported to target TNNT2 and TNNI3. Using the best performing protocol (Protocol 2), the best performing antibody clones (anti-TNNT2 clones 1C11 and 13-11, anti-TNNI3 clones 2Q1100 and C5) were titrated to determine the appropriate amount of antibody to use for flow cytometry. Titration is a quintessential step for any antibody-based technique and is especially critical for flow cytometry. Notably different amounts of anti-TNNI3 were optimal for the two tested clones (0.5μg for clone 2Q1100 vs 0.1μg for clone C5) despite the fact they are reported to target the same epitope on the same protein.

Using the optimal antibody-to-cell ratio as determined by the titration assay, an epitope competition assay using synthetic peptides was used to test the specificity of the antibodies for their reported epitopes. Only one of the three antibodies, anti-TNNI3 clone 2Q1100, was significantly blocked from binding to cells by pre-incubation with its peptide antigen despite sharing this epitope with anti-TNNI3 clone C5. Notably, the homologous peptides from TNNI1 and TNNI2 were also able to block signal, although this may be an artifact of the 10,000x molar equivalents used to observe blocking. Overall, these experiments were unable to provide conclusive evidence that the three antibody clones tested were specific for their reported peptide antigens. However, the inability of the linear peptides to block the antibodies could be indicative of either specificity for a different epitope or reliance on a secondary or tertiary structure that is absent outside of the context of the protein. Further challenges to interpreting these data relate to the difficulty in obtaining technical details regarding how commercially available antibody epitopes were originally mapped. Though these experiments do not provide evidence to support the specificity of these antibodies to their stated epitopes, neither do they preclude the specificity of these antibodies for their reported protein targets. It remains possible that multiple isoforms of TNNI are detected by the anti-TNNI3 clones used here. Based on the high degree of sequence homology among TNNI isoforms, it is possible that an anti-TNNI3 antibody that targets the N-terminal extension of TNNI3 would be best able to discriminate among TNNI isoforms. Although clones EP1106Y and 4C2 are reported to target this unique sequence, we did not find that these antibodies provided suitable results for the goals of this study. Nonetheless, the results of the co-immunodetection experiment and tests on cardiac fibroblasts suggest the best performing antibodies (2Q1100, C5, 1C11) are specifically detecting cardiomyocytes in this context. Finally, the mixed population experiments demonstrate the capacity of the antibodies and protocol to discriminate between positive and negative cells within the same experimental sample, which is the ultimate test of suitability for assessing population heterogeneity within hPSC-CM cultures. As stated previously and discussed in detail below, antibody and protocol suitability is context dependent. Therefore, despite the cell-type specificity demonstrated in this study, further validation would be required to demonstrate whether these antibodies and methods could distinguish between cardiomyocytes and other cell types (*e.g*. skeletal or smooth muscle cells) within other contexts (*e.g*. imaging). The SOP developed here was evaluated for its ability to accurately assess heterogeneity when applied to different cell lines and differentiation protocols in two additional laboratories, further establishing its reliability and reproducibility.

The accurate execution of a flow cytometry experiment requires attention to many technical details. Unfortunately, in literature published in the past seven years for hPSC-CM, >26% and >15% studies failed to report detailed sample preparation conditions and antibody clone information, respectively. Importantly, as demonstrated by the results in Figure 3, fixation and permeabilization conditions can drastically affect the measured flow cytometry signal, an effect that is clone-dependent. For example, the signal for anti-TNNT2 clone 13-11, the most commonly used antibody in published literature, is highly sensitive to permeabilization conditions. As shown in Figure 3, when using a methanol-based permeabilization (Protocol 1), this antibody shows significant overlap of the negative and positive populations within the histogram. Among the published studies surveyed here, five different permeabilization conditions ranging in reagent concentrations (including 11% that used methanol), were used with this antibody clone. As the signal obtained in a flow cytometry experiment is also dependent on additional variables, including incubation time and blocking solution composition, it is not possible to definitively conclude that data from such studies are problematic. However, as these details are often not reported or underestimated, we believe it is prudent to highlight this antibody’s sensitivity to permeabilization in light of its popularity. Importantly, the effect of sample preparation conditions on antibody utility is cell-type dependent, meaning that results for a single antibody clone can vary among cell types when different sample preparation strategies are applied. For example, consider the data for anti-TNNI3 clone 19C7. Specifically, the intensity of the hPSC and hPSC-CM histograms change, and they overlap to a varying degree based on sample preparation protocol. Overall, these data clearly demonstrate how technical details can drastically affect results obtained and highlight why all experimental variables must be empirically tested on individual clones and cell types, as performance of one antibody is not predictive for another antibody under the same conditions. Also, these data highlight why simply comparing signals from an antibody to that of an equivalent amount of isotype control is insufficient to conclude an antibody is detecting the desired target, hence why negative cell type controls are essential. Based on the literature survey, data from negative cell type controls are not typically reported, so it is unclear whether they are routinely implemented and not reported or rather not included. In addition to antibody clone selection, cell collection, and sample preparation, another important aspect that requires attention to detail includes data acquisition and analysis. Flow cytometry data are dependent on instrument characteristics (*e.g*. laser strength, filter block selection, and detector sensitivity). Therefore, recording these experimental details, as advised (Lee et al., 2008), is a suggestion we enthusiastically echo. Once data are collected, they must be analyzed, and this is a step that can introduce bias and influence interpretation. Consequently, important considerations for data collection and analysis, including suggestions regarding the number of events to collect and how to adjust laser power settings, are provided in our SOP. Finally, considering the observed lack of details reported in published literature, our experimental observations regarding key experimental details that drastically affect data quality, and recent calls for data reporting guidelines and standards (Lee et al., 2008), we have generated a suggested “Checklist for Publication” in the SOP which contains details to be included when publishing flow cytometry based assessments of hPSC-CM, including sample preparation information and controls that are important for enabling the successful replication and interpretation of experimental data.

Although these studies focused on the use of antibodies to detect intracellular markers, alternative strategies are possible. For example, genetic-modification of cell lines to express a transgene marker (*e.g*. GFP) driven by a cell-type or tissue specific promoter can offer a convenient antibody-independent strategy to assess heterogeneity within a cell population. However, it is not always practical to generate transgenic lines, especially for high-throughput studies of hiPSC derived from multiple patients. Although cell surface markers offer the significant advantage of being amenable to detection on live cells, thereby enabling live cell sorting, a single cardiomyocyte-specific surface marker has not yet been widely validated, although there are reports of marker combinations (Skelton et al., 2014) and markers of cardiomyogenic progenitors (Dubois et al., 2011; Yang et al., 2008) that can be helpful in assessing cell identity. Moreover, detection of cell surface proteins can be complicated because of their potential sensitivity to the enzymatic conditions necessary to prepare single-cell suspensions for flow cytometry, and the biological effects that can be triggered by the binding of antibodies to critical cell surface proteins. For these reasons, the current study focused on developing an SOP that is universally applicable to high-throughput hiPSC studies and uses well validated markers for cardiomyocyte identity. Immunofluorescent microscopy is another technique that can be used for quality assessment of hPSC-CM differentiation cultures. Microscopy offers the ability to visualize the localization of an analyte, but quantitation of cellular heterogeneity or antigen abundance is challenging. In this way, flow cytometry is advantageous as it provides quantitative, single-cell measurements to accurately assess population heterogeneity with high sensitivity. However, it can be difficult to dissociate adherent cells and maintain cellular integrity during sample processing. Therefore, it is possible that different cell types (even from the same well) respond differently to collection and sample preparation strategies. As such, perhaps the most critical step for accurate flow cytometry data is the collection of cells and the preparation of a single-cell suspension of viable cells. Differences in cell dissociation and associated cellular integrity can serve as confounding variables when making inferences regarding the population based on the measured sample (*i.e*. the cells that make it into the cytometer). Therefore, imaging remains an important complement to flow cytometry.

In conclusion, while flow cytometry offers the advantages of high-throughput, population-based, single-cell and quantitative analyses, accuracy of measurement is dependent on numerous technical variables which are often overlooked and under-reported. To facilitate enhanced rigor regarding the application of flow cytometry for the assessment of heterogeneity within hPSC-CM cultures, a comprehensive SOP based on the results of the current study is provided. The SOP contains the detailed experimental protocol and a substantial number of observations, suggestions, and considerations for customization to assist new users in its implementation. Of course, we advocate that laboratories validate this protocol independently for their own cell lines and differentiation protocols. However, as the current study demonstrates its replicability among three laboratories, the SOP is expected to benefit both established laboratories and those new to this field. Finally, we present a workflow for establishing the fit-for-purpose use of other antibody clones or protocols in contexts beyond the assessment of hPSC-CM. Overall, we hope that adhering to rigorous standards for antibody validation and use, reporting of experimental details, and presentation of data, these studies will promote enhanced utility and dialogue regarding hPSC-CM for a variety of research and translational applications.

## Experimental Procedures

A complete record of experimental details for all procedures can be found in Supplemental Information.

### Cell Culture and Reagents

Laboratory 1 (used for initial screen and all method development) - DF6-9-9T hiPSCs were maintained in monolayer culture and differentiation performed as described (Bhattacharya et al., 2014; Kropp et al., 2015). Laboratory 2 – hPSC-CM were generated from hPSC line 19-3 from a healthy donor. Pluripotent cells were differentiated to cardiomyocytes as described (Burridge et al., 2015). Laboratory 3 – Undifferentiated hESCs (H7) were induced to differentiate as described (Wang et al., 2015). All experiments were performed using hPSC-CM from days 20-25 of differentiation. Normal human ventricular cardiac fibroblasts (Lonza Cat # CC-2904) were cultured per vendor’s recommendations.

### Parallel Reaction Monitoring

Day 25 hPSC-CM cell lysate and recombinant human TNNI3 (ProSpec, PRO-324) were digested with trypsin and analyzed by liquid chromatography tandem mass spectrometry (LC-MS/MS) using an Orbitrap Fusion Lumos (Thermo). In initial discovery studies, data dependent acquisition was used for selection of peptides which had favorable characteristics (*i.e*. well-defined chromatographic peak, easily ionized). Subsequently, stable isotopically labeled synthetic peptides were obtained for the three best-performing TNNI3 peptides and used as internal standards. A targeted mass spectrometry assay was developed using parallel reaction monitoring (Peterson et al., 2012) to selectively detect three peptides from TNNI3 and four peptides from TNNT2 at their observed mass and time of elution from preliminary experiments of recombinant protein and cell lysate, respectively. This assay was applied to day 25 hPSC-CMs and hPSC and cardiac tissue samples were included as negative and positive controls, respectively.

### Flow cytometry

Three protocols were used to prepare hPSC-CM for flow cytometry (Table 1). All protocols were performed at room temperature using 100 μL for all solutions unless otherwise indicated. For the initial screen, the amount of antibody used was selected based on published literature or manufacturer’s recommendation when available. When unavailable, 1 μg was used. The initial screen was performed using two biological replicates, while all subsequent analyses were performed using three biological replicates. Data were acquired on a BD LSR II flow cytometer using the filter cubes described in Table 1 and were analyzed using FlowJo v.10 (FlowJo LLC).

### Epitope Competition Assay

Synthetic peptides (>99% purity, Genscript) with sequences representing the epitopes included TNNI1 (VEVGDWR), TNNI2 (RDVGDWR), TNNI3 (REVGDWR) and TNNT2 (EEEENRRKAEDEARKKKALSN) were generated and resuspended according to manufacturer’s guidelines. Respective antibodies were incubated with each blocking peptide (10,000x molar excess of antibody) for 30 min at room temperature in blocking solution. Following incubation, this peptide-antibody mix was added to the fixed cell sample and sample preparation proceeded according to protocol 2 (Table 1).

### Co-Immunodetection Experiments

For co-immunodetection, primary antibodies were added simultaneously in two combinations: anti-TNNT2 clone 1C11/anti-TNNI3 clone 2Q1100 and anti-TNNT2 clone 1C11/anti-TNNI3 clone C5. To avoid the complication of spectral overlap, the secondary antibody used in co-immunodetection experiments for anti-TNNI3 clone C5 was anti-mouse IgG2b-AlexaFluor647 (Thermofisher Cat. # A21242).

### Cell Mixing Experiments

Samples were generated by mixing hPSC-CM with hPSC at ratios of 100:0, 75:25, 50:50, 25:75 and 0:100 such that the total number of cells in the mixed sample was 1×10^6^ and subsequently prepared according to protocol 2 (Table 1). A finalized protocol (Supplemental Information) was shared with two laboratories (Dr. Paul Burridge, Northwestern; Dr. Kenneth Boheler, Hong Kong University), each of which used different cell lines and differentiation protocols than used in SOP development.

## Author Contributions

R.L.G. and M.W. conceived the study; M.W., R.W., and R.L.G. designed experiments and analyzed the data; R.L.G. supervised the study; M.W., R.W., and R.L.G. co-wrote the manuscript; M.W. and R.W. performed flow cytometry experiments; M.W. performed mass spectrometry experiments; E.K. performed immunostaining; M.W., R.W., and E.K. wrote the standard operating protocol; E.P., K.R.B., M.R. and P.W.B. performed cell mixing experiments to assess the protocol. All authors approved the final manuscript.

## Acknowledgements

This work was supported by the National Institutes of Health [R01-HL126785 and R01-HL134010 to RLG; F31-HL140914 to MW; R00-HL121177 and R01-CA2200002 to PWB]; American Heart Association [15GRNT24980002 to RLG and 18TPA34230105 to PWB] and Entopsis Asia, Ltd to KRB. EMK is a member of the MCW-MSTP which is partially supported by a T32 grant from NIGMS, GM080202. Funding sources were not involved in study design, data collection, interpretation, analysis or publication. We thank Dr. Paul Goldspink, Dr. Christopher Ashwood, and Linda Berg Luecke (MCW) for insightful discussions and careful review of the manuscript. Mass spectrometry analyses were performed using instrumentation in the Center for Biomedical Mass Spectrometry Research at the Medical College of Wisconsin. Flow cytometry analyses were performed using instrumentation in the Blood Center of Wisconsin Flow Cytometry Core.

## References

Addis, R.C., Ifkovits, J.L., Pinto, F., Kellam, L.D., Esteso, P., Rentschler, S., Christoforou, N., Epstein, J.A., and Gearhart, J.D. (2013). Optimization of direct fibroblast reprogramming to cardiomyocytes using calcium activity as a functional measure of success. J Mol Cell Cardiol 60, 97–106.

Aebersold, R., Burlingame, A.L., and Bradshaw, R.A. (2013). Western blots versus selected reaction monitoring assays: time to turn the tables? Mol Cell Proteomics 12, 2381–2382.

Anderson, P.A., Malouf, N.N., Oakeley, A.E., Pagani, E.D., and Allen, P.D. (1991). Troponin T isoform expression in humans. A comparison among normal and failing adult heart, fetal heart, and adult and fetal skeletal muscle. Circ Res 69, 1226–1233.

Batalov, I., and Feinberg, A.W. (2015). Differentiation of Cardiomyocytes from Human Pluripotent Stem Cells Using Monolayer Culture. Biomark Insights 10, 71–76.

Bedada, F.B., Chan, S.S., Metzger, S.K., Zhang, L., Zhang, J., Garry, D.J., Kamp, T.J., Kyba, M., and Metzger, J.M. (2014). Acquisition of a quantitative, stoichiometrically conserved ratiometric marker of maturation status in stem cell-derived cardiac myocytes. Stem Cell Reports 3, 594–605.

Bhattacharya, S., Burridge, P.W., Kropp, E.M., Chuppa, S.L., Kwok, W.M., Wu, J.C., Boheler, K.R., and Gundry, R.L. (2014). High efficiency differentiation of human pluripotent stem cells to cardiomyocytes and characterization by flow cytometry. J Vis Exp, 52010.

Bhavsar, P.K., Dhoot, G.K., Cumming, D.V., Butler-Browne, G.S., Yacoub, M.H., and Barton, P.J. (1991). Developmental expression of troponin I isoforms in fetal human heart. FEBS Lett 292, 5–8.

Bicer, S., and Reiser, P.J. (2013). Complex tropomyosin and troponin T isoform expression patterns in orbital and global fibers of adult dog and rat extraocular muscles. J Muscle Res Cell Motil 34, 211–231.

Bodor, G.S., Porterfield, D., Voss, E.M., Smith, S., and Apple, F.S. (1995). Cardiac troponin-I is not expressed in fetal and healthy or diseased adult human skeletal muscle tissue. Clin Chem 41, 1710–1715.

Bodor, G.S., Survant, L., Voss, E.M., Smith, S., Porterfield, D., and Apple, F.S. (1997). Cardiac troponin T composition in normal and regenerating human skeletal muscle. Clin Chem 43, 476–484.

Bordeaux, J., Welsh, A., Agarwal, S., Killiam, E., Baquero, M., Hanna, J., Anagnostou, V., and Rimm, D. (2010). Antibody validation. Biotechniques 48, 197–209.

Bradbury, A., and Pluckthun, A. (2015). Reproducibility: Standardize antibodies used in research. Nature 518, 27–29.

Brooks, H.L., and Lindsey, M.L. (2018). Guidelines for Authors and Reviewers on Antibody Use in Physiology Studies. Am J Physiol Heart Circ Physiol.

Burridge, P.W., Matsa, E., Shukla, P., Lin, Z.C., Churko, J.M., Ebert, A.D., Lan, F., Diecke, S., Huber, B., Mordwinkin, N.M., et al. (2014). Chemically defined generation of human cardiomyocytes. Nat Methods 11, 855–860.

Dubois, N.C., Craft, A.M., Sharma, P., Elliott, D.A., Stanley, E.G., Elefanty, A.G., Gramolini, A., and Keller, G. (2011). SIRPA is a specific cell-surface marker for isolating cardiomyocytes derived from human pluripotent stem cells. Nat Biotechnol 29, 1011–1018.

Ebashi, S. (1983). Regulation of muscle contraction. Cell Muscle Motil 3, 79–87.

Fu, J.D., Stone, N.R., Liu, L., Spencer, C.I., Qian, L., Hayashi, Y., Delgado-Olguin, P., Ding, S., Bruneau, B.G., and Srivastava, D. (2013). Direct reprogramming of human fibroblasts toward a cardiomyocyte-like state. Stem Cell Reports 1, 235–247.

Hunkeler, N.M., Kullman, J., and Murphy, A.M. (1991). Troponin I isoform expression in human heart. Circ Res 69, 1409–1414.

Kajioka, S., Takahashi-Yanaga, F., Shahab, N., Onimaru, M., Matsuda, M., Takahashi, R., Asano, H., Morita, H., Morimoto, S., Yonemitsu, Y., et al. (2012). Endogenous cardiac troponin T modulates Ca(2+)-mediated smooth muscle contraction. Sci Rep 2, 979.

Kropp, E.M., Oleson, B.J., Broniowska, K.A., Bhattacharya, S., Chadwick, A.C., Diers, A.R., Hu, Q., Sahoo, D., Hogg, N., Boheler, K.R., et al. (2015). Inhibition of an NAD(+) salvage pathway provides efficient and selective toxicity to human pluripotent stem cells. Stem Cells Transl Med 4, 483–493.

Lee, J.A., Spidlen, J., Boyce, K., Cai, J., Crosbie, N., Dalphin, M., Furlong, J., Gasparetto, M., Goldberg, M., Goralczyk, E.M., et al. (2008). MIFlowCyt: the minimum information about a Flow Cytometry Experiment. Cytometry A 73, 926–930.

Lian, X., Bao, X., Zilberter, M., Westman, M., Fisahn, A., Hsiao, C., Hazeltine, L.B., Dunn, K.K., Kamp, T.J., and Palecek, S.P. (2015). Chemically defined, albumin-free human cardiomyocyte generation. Nat Methods 12, 595–596.

Mummery, C.L., Zhang, J., Ng, E.S., Elliott, D.A., Elefanty, A.G., and Kamp, T.J. (2012). Differentiation of human embryonic stem cells and induced pluripotent stem cells to cardiomyocytes: a methods overview. Circ Res 111, 344–358.

Ohno, Y., Yuasa, S., Egashira, T., Seki, T., Hashimoto, H., Tohyama, S., Saito, Y., Kunitomi, A., Shimoji, K., Onizuka, T., et al. (2013). Distinct iPS Cells Show Different Cardiac Differentiation Efficiency. Stem Cells Int 2013, 659739.

Peterson, A.C., Russell, J.D., Bailey, D.J., Westphall, M.S., and Coon, J.J. (2012). Parallel reaction monitoring for high resolution and high mass accuracy quantitative, targeted proteomics. Mol Cell Proteomics 11, 1475–1488.

Puppala, D., Collis, L.P., Sun, S.Z., Bonato, V., Chen, X., Anson, B., Pletcher, M., Fermini, B., and Engle, S.J. (2013). Comparative gene expression profiling in human-induced pluripotent stem cell--derived cardiocytes and human and cynomolgus heart tissue. Toxicol Sci 131, 292–301.

Rittoo, D., Jones, A., Lecky, B., and Neithercut, D. (2014). Elevation of cardiac troponin T, but not cardiac troponin I, in patients with neuromuscular diseases: implications for the diagnosis of myocardial infarction. J Am Coll Cardiol 63, 2411–2420.

Sasse, S., Brand, N.J., Kyprianou, P., Dhoot, G.K., Wade, R., Arai, M., Periasamy, M., Yacoub, M.H., and Barton, P.J. (1993). Troponin I gene expression during human cardiac development and in end-stage heart failure. Circ Res 72, 932–938.

Sheng, J.J., and Jin, J.P. (2016). TNNI1, TNNI2 and TNNI3: Evolution, regulation, and protein structure-function relationships. Gene 576, 385–394.

Skelton, R.J., Costa, M., Anderson, D.J., Bruveris, F., Finnin, B.W., Koutsis, K., Arasaratnam, D., White, A.J., Rafii, A., Ng, E.S., et al. (2014). SIRPA, VCAM1 and CD34 identify discrete lineages during early human cardiovascular development. Stem Cell Res 13, 172–179.

Thavandiran, N., Dubois, N., Mikryukov, A., Masse, S., Beca, B., Simmons, C.A., Deshpande, V.S., McGarry, J.P., Chen, C.S., Nanthakumar, K., et al. (2013). Design and formulation of functional pluripotent stem cell-derived cardiac microtissues. Proc Natl Acad Sci U S A 110, E4698–4707.

Tompkins, J.D., Jung, M., Chen, C.Y., Lin, Z., Ye, J., Godatha, S., Lizhar, E., Wu, X., Hsu, D., Couture, L.A., et al. (2016). Mapping Human Pluripotent-to-Cardiomyocyte Differentiation: Methylomes, Transcriptomes, and Exon DNA Methylation “Memories”. EBioMedicine 4, 74–85.

Wang, Y., Li, Z.C., Zhang, P., Poon, E., Kong, C.W., Boheler, K.R., Huang, Y., Li, R.A., and Yao, X. (2015). Nitric Oxide-cGMP-PKG Pathway Acts on Orai1 to Inhibit the Hypertrophy of Human Embryonic Stem Cell-Derived Cardiomyocytes. Stem Cells 33, 2973–2984.

Yang, L., Soonpaa, M.H., Adler, E.D., Roepke, T.K., Kattman, S.J., Kennedy, M., Henckaerts, E., Bonham, K., Abbott, G.W., Linden, R.M., et al. (2008). Human cardiovascular progenitor cells develop from a KDR+ embryonic-stem-cell-derived population. Nature 453, 524–528.

